# The novel linkage between *Fuz* and *Gpr161* genes regulates sonic hedgehog signaling during mouse embryonic development

**DOI:** 10.1101/2024.01.11.575263

**Authors:** Sung-Eun Kim, Hyun Yi Kim, Bogdan J. Wlodarczyk, Richard H. Finnell

## Abstract

Sonic hedgehog (Shh) signaling regulates embryonic morphogenesis utilizing primary cilia, the cell antenna acting as a signaling hub. Fuz, an effector of planar cell polarity (PCP) signaling, involves Shh signaling via cilia formation, while the G protein-coupled receptor 161 (Gpr161) is a negative regulator of Shh signaling. The range of phenotypic malformations observed in mice bearing mutations in either of these two genes is similar; however, their functional relations have not been previously explored. This study identified the genetic and biochemical link between Fuz and Gpr161 in mouse embryonic development. *Fuz* was genetically epistatic to *Gpr161* via Shh signaling during mouse embryonic development. The FUZ biochemically interacted with GPR161, and Fuz regulated Gpr161 ciliary trafficking via β-arrestin2. Our study suggested the novel Gpr161-Fuz axis that regulates Shh signaling during mouse embryonic development.

**Summary statement:** This study illuminates the novel genetic and biochemical linkages between Fuz and Gpr161 to regulate sonic hedgehog signaling during mouse embryonic development.

## Introduction

Sonic hedgehog (Shh) signaling is one of the most critical morphogen signaling pathways for embryonic morphogenesis and organogenesis (Matise and Wang, 2011; Alvarez-Buylla and Ihrie, 2014; Briscoe and Small, 2015; Xavier et al., 2016). The morphogen gradients of Shh and others, such as Wnt and Bone morphogenetic protein (BMP), often lead to the spatiotemporal regulation of gene expression to determine the cell fates essential for the proper tissue patterning (Groves et al., 2020). Specifically, the Shh morphogen is secreted from the floor plate and/or notochord and determines the ventral side of the neural tube. The gradient of Shh morphogen counterbalances with that of Wnt or BMP morphogens, secreted from the roof plate of the neural tube, to generate the proper neural tube patterning for determining the neuronal cell identities (Briscoe and Small, 2015). Shh signaling depends on the primary cilia as many signaling proteins in Shh signaling are trafficking within primary cilia responding to the extracellular stimuli (Sasai and Briscoe, 2012). Primary cilia are antenna-like cellular protrusive organelles essential for multiple cellular signaling pathways for embryonic development and homeostasis (Goetz and Anderson, 2010). The microtubule-based transporting systems, so-called intraflagellar transports (IFTs), involve anterograde and retrograde movements of signaling proteins within primary cilia, coupled with a kinesin or dynein motor protein, respectively (Mill et al., 2023). In addition, IFTs are involved in ciliogenesis by recruiting and transporting required proteins to the basal body of primary cilia. At present, the mechanisms and functions of Shh signaling protein trafficking within cilia and its effect on Shh signaling are not fully understood.

Fuzzy (Fuz) is considered a member of the CPLANE (ciliogenesis and planar polarity effectors) complex, including Intu, Fuz, and Wdpcp proteins (Langousis et al., 2022) and serves as an effector of planar cell polarity (PCP) signaling (Gray et al., 2009). Fuz contributes to primary cilia formation via delivering Rab8 and Disheveled (Dvl) into the basal body (Zilber et al., 2013) and retrograde protein trafficking in the primary cilia. The genetic ablation of *Fuz* in Xenopus creates convergent extension and cilia defects (Park et al., 2006). The *Fuz* mutant mice showed milder abnormal phenotype involving defects in PCP signaling (i.e., kinked tail and cardiac defects) and Shh signaling potentially secondary to compromised ciliogenesis (i.e., cranial neural tube defects (NTDs), polydactyly and skeletal defects) without convergent extension defects (Park et al., 2006; Gray et al., 2009). In addition, this locus has been implicated as a genetic risk factor in human neural tube defects (Seo et al., 2011) and craniosynostosis (Barrell et al., 2022). Heydeck et al (Heydeck et al., 2009) showed that *Fuz* mutant mice presented with gene expression changes in the ventral neural tubes and reduced Shh signaling in forelimb buds. In addition, the cilia numbers and lengths were substantially reduced in limb buds and the spinal neural tube in *Fuz* mutant mice. Taken together, Fuz regulates Shh signaling potentially via ciliogenesis in embryonic limb and neural tube development. Yet, the molecular mechanism by which Fuz regulates Shh signaling and its signaling partners has not been adequately characterized.

G protein-coupled receptor 161 (Gpr161) is a negative regulator of Shh signaling, and its inhibitory activity is maintained by its localization in the primary cilia (Mukhopadhyay et al., 2013). Without Shh signal, Gpr161 is localized in the primary cilia and activates protein kinase A (PKA) subsequent to the increased cAMP levels via Gαs. The activated PKA proteolyzed Gli3 full length (Gli3-FL) into Gli3 repressor form (Gli3-R), thereby repressing Shh-responsive transcription. In the presence of the Shh signal, Gpr161 is removed from the primary cilia via recruitment of β-arrestin2 and Grk2 (Pal et al., 2015), and its repressor activities on Shh signaling are retained. Subsequently, Shh responsive transcription was turned on, and the expression of Shh signaling target genes, such as Gli1 and Ptch1, that involve multiple morphogenetic processes were elevated. For the ciliary localization of GPR161, IFT-A complex, and TULP3 coordinate the ciliary entry of GPR161 (Hirano et al., 2017), and BBSome mediates the removal of GPR161 from the primary cilia (Nozaki et al., 2018; Ye et al., 2018). Specifically, IFT-A complex involves both entry of GPR161 into the cilia and the retrograde trafficking within the cilia. The *GPR161* allele is associated with the etiology of NTDs in humans and mice (Kim et al., 2019) and is additionally associated with craniofacial defects and limb abnormalities in mouse models (Hwang et al., 2018; Hwang et al., 2021; Kim et al., 2021; Kim et al., 2023).

The range of phenotypic malformations in *Fuz* and *Gpr161* mutant mice is similar, including NTDs, craniofacial defects, and limb defects, and we hypothesized that *Fuz* and *Gpr161* are genetically related during mouse embryonic development. In this study, we investigated the molecular and genetic relations of *Fuz* and *Gpr161* during mouse embryonic development and in regulating Shh signaling. We have demonstrated that *Fuz* was genetically epistatic to *Gpr161* during mouse embryonic spinal neural tube development via Shh signaling regulation. We further observed the biochemical interaction between FUZ and GPR161 and the role of FUZ in the ciliary trafficking of GPR161 in the primary cilia. Our results illuminate the novel genetic and biochemical linkage between *Fuz* and *Gpr161* in regulating Shh signaling and its role in mouse embryonic development.

## Results

### *Fuz* is genetically epistatic to *Gpr161* in mouse spinal neural tube development

The *Fuz* and *Gpr161* null embryos share strikingly similar phenotypic malformations regarding rostral/spinal NTDs and craniofacial defects (Gray et al., 2009; Kim et al., 2019; Kim et al., 2021). We questioned a possible genetic connection that might exist during mouse embryonic development. To answer this question, we generated *Fuz; Gpr161* double knockout (DKO) (*Fuz-/-; Gpr161-/-*, hereafter DKO) mice to compare its gross phenotype with *Fuz* or *Gpr161* single knockout (KO) embryos. The *Fuz; Gpr161* DKO mice were embryonic lethal by E12.5, and the compound heterozygotes (*Fuz+/-; Gpr161+/-*) were viable without obvious phenotypic malformations. As previously reported, *Gpr161* KO (*Gpr161-/-*) embryos presented with severe rostral and spinal NTDs often together with craniofacial defects with 100% penetrance (Mukhopadhyay et al., 2013; Kim et al., 2019), and *Fuz* KO (*Fuz-/-*) embryos showed rostral NTDs with craniofacial defects and kinked tails with about 60% penetrance (Gray et al., 2009). We observed no phenotypic abnormalities in the spinal neural tube of DKO embryos (Figure 1A: dotted lines) and less severe malformations in craniofacial and rostral NTDs (Figure 1A: arrows and arrowheads) in DKO embryos at E9.5 and E10.5 compared to *Gpr161* single KO embryos (Figure 1A). In addition, DKO embryos survived longer (by E12.5) than *Gpr161* single KO embryos (by E10.5), but shorter than *Fuz* single KO fetuses (by E18.5).

**Figure 1.**
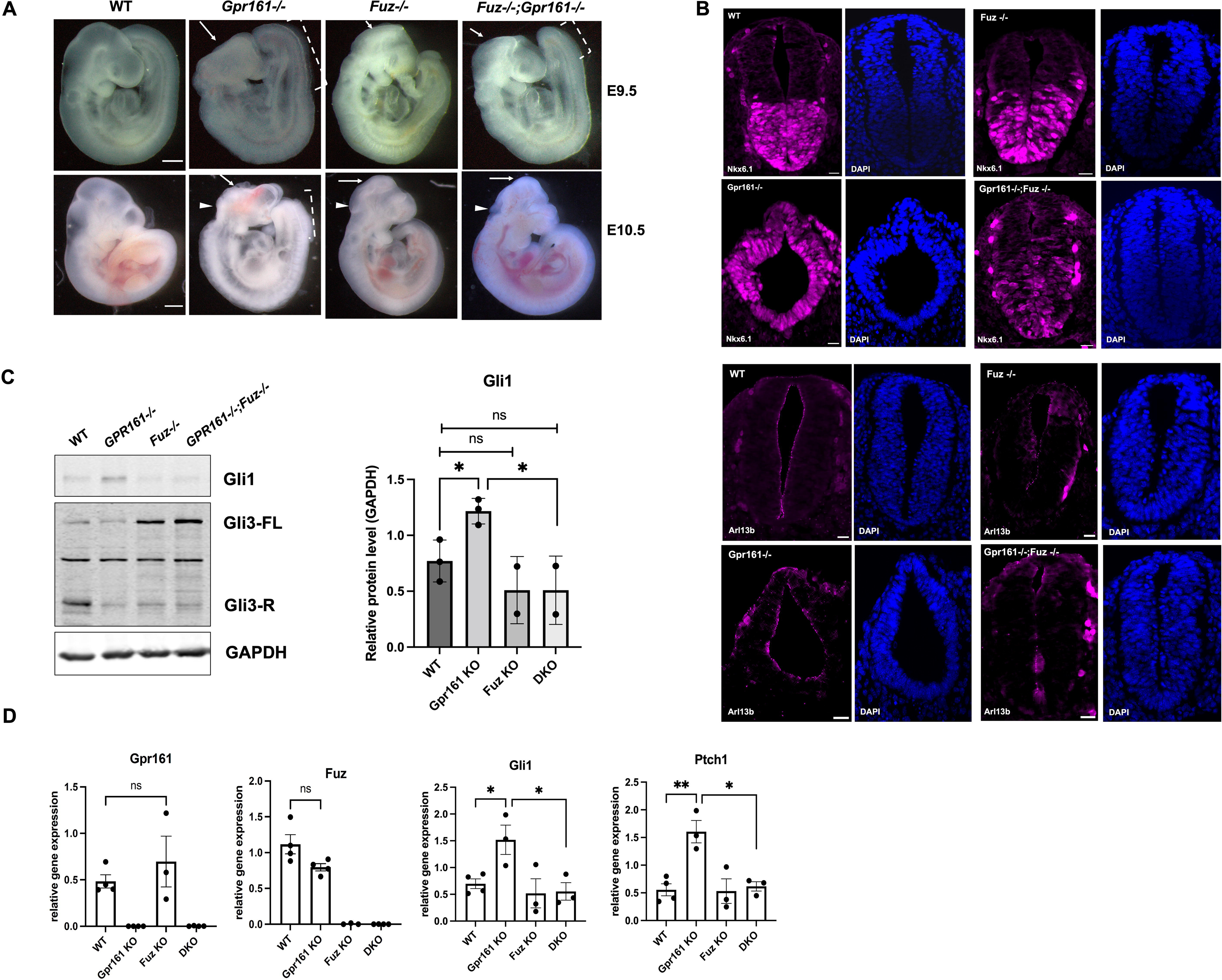
The gross phenotypic malformation, neural tube patterning, primary cilia and the status of Shh signaling activation of WT, *Fuz* KO, *Gpr161* KO and *Gpr161:Fuz* double knock out (DKO) embryos at E9.5 and E10.5. (A) The gross phenotypic malformation in WT, *Fuz* KO, *Gpr161* KO, and DKO embryos at E9.5 and E10.5. Dotted line: open PNP length, Arrow: midbrain NTs, and Arrowhead: hindbrain NTs. Scale bars: 1 mm. (B) The immunostaining of Nkx6.1 and Arl13b in the spinal neural tube of WT, *Fuz* KO, *Gpr161* KO, and DKO embryos at E10.5 (n=3 for each genotyped embryo). Scale bars: 20 μm. (C) The western blot analysis with the lysates from the whole embryos of WT, *Fuz* KO, *Gpr161* KO, and DKO at E9.5. Gli1, Gli3 and Gapdh proteins were detected. (WT: n=3, *Fuz* KO: n=2, *Gpr161* KO: n=3, DKO: n=2), Right: The Gli1 protein intensities were quantified and normalized to Gapdh. (*<0.05) (D) The qRT-PCR with the whole embryos of WT, *Fuz* KO, *Gpr161* KO, and DKO at E9.5. The expression of *Gpr161*, *Fuz*, *Gli1*, and *Ptch1* was normalized to *Gapdh*. (WT: n=4, *Fuz* KO: n=3, *Gpr161* KO: n=4, DKO: n=4) (*<0.05, **<0.005)

As DKO embryos showed fully rescued spinal NTDs compared to *Gpr161* single KO embryos, we observed the status of neural tube patterning in the spinal neural tube of DKO embryos and compared it to *Gpr161* and *Fuz* single KO embryos. Nkx6.1 is mainly expressed in the V2 interneurons and motor neurons (MN) in the ventral side of the neural tube (Sander et al., 2000). As shown in the previous study (Mukhopadhyay et al., 2013), Nkx6.1 expression in *Gpr161* KO embryos was expanded to the dorsal side of the neural tube compared to that in WT embryos (Figure 1B: Upper). On the other hand, the Nkx6.1 expression pattern of WT, *Fuz* KO, and DKO embryos was similar in the ventral aspect. The primary cilia are present in the apical side of neural tubes, and Fuz is involved in the cilliogenesis (Gray et al., 2009; Vogel et al., 2012). Hence, we examined the presence of primary cilia in the neural tube of each genotyped embryo (Figure 1B: Lower). The numbers of cilia were substantially reduced in the spinal neural tube of *Fuz* KO and DKO embryos, while those in *Gpr161* KO embryos were as intact as those of WT embryos. These results suggest that the restored ventralized neural tube patterning and reduced numbers of primary cilia in DKO embryos might be related to the rescued spinal NTDs in DKO embryos compared to those in *Gpr161* KO embryos.

As Shh signaling regulates the neural tube patterning (Tanabe and Jessell, 1996), we compared the activation status of Shh signaling in WT, *Fuz* KO, *Gpr161* KO and DKO embryos (Figures 1C and 1D). The Gli1 protein expression in DKO embryos was similar to that of WT and *Fuz* KO embryos, while the Gli1 protein level was elevated in *Gpr161* KO embryos (Figure 1C). Consistently, the patterns of *Gli1* and *Ptch1* gene expression, which are Shh target genes, were similar to the pattern of Gli1 protein expression (Figure 1D). These results suggest that the increased Shh signaling in *Gpr161* KO embryos was reduced in DKO embryos, which could explain the restored ventralized patterning observed in DKO embryos.

### The differential gene expression profiles of the anterior and posterior regions of embryos

We performed RNA sequencing (RNA seq) experiments to explore the unbiased gene expression profile changes in the embryos of each genotype (WT, *Gpr161* KO, *Fuz* KO and DKO). The spinal NTDs seen consistently in *Gpr161* KO embryos were fully rescued in DKO embryos, whereas craniofacial defects and cranial NTDs in *Gpr161* KO embryos were only partially rescued in DKO embryos (Figure 1A). Therefore, we collected the anterior and posterior regions of WT, *Fuz* KO, *Gpr161* KO, and DKO embryos separately to compare the differential gene expression patterns between each embryo’s rostral and spinal regions. The principal component analysis (PCA) and volcano plots represented (Supplementary Figures 1 and 2) the severity and prevalence of phenotypic malformations of each genotype and region of embryos (Supplementary Figures 1 and 2). We selected twenty of the most differentially expressed genes (DEGs) from each embryo group’s anterior and posterior regions. Shh signaling-related genes were included in the top 20 DEGs in both anterior (*Foxl1*, *Shh*, *Nkx2-4*, and *Gpr161*) (Figure 2B) and posterior (*Nkx2-2*, *Nkx2-9*, *Ferd3l*, and *Gpr161*) regions of embryos (Figure 2D). The overlapping DEGs were not observed between the anterior and posterior regions of the embryos, except *Gpr161* and *Fuz*, suggesting the differential gene sets contributed to the head and tail morphogenesis. Among the top 20 DEGs in the anterior portions of the embryos, *Shh*, *Tyr*, *Tryp1*, and *Aldh1a1* genes (Tief et al., 1997; Anthony and Heintz, 2007) were relevant to neural tube development, and *Six6* and *Jhy* genes (Muniz-Talavera and Schmidt, 2017; Diacou et al., 2022) were related to eye and brain development, respectively. In the top 20 DEGs in the tails of embryos, *Nkx2-2*, *Nkx2-9*, *Ferd3l*, *Gpr50*, and *Atoh1* genes (Pabst et al., 1998; Briscoe et al., 2000; Ebert et al., 2003; Mansour et al., 2014; Azzara et al., 2021), along with *Gpr161* and *Fuz*, were relevant to spinal neural tube development. We observed three gene clusters where DEGs were highly enriched and performed a Gene Ontology (GO) analysis. In anterior region of the embryos, genes involved in forebrain development, cell fate commitment, pattern specification, regionalization, neuron differentiation, muscle differentiation, and heart morphogenesis were enriched in three clusters (Figure 2C). In the posterior region of the embryos, genes involved in extracellular structure/matrix organization, epithelial tube morphogenesis, cell fate commitment, and muscle organ development were enriched in three clusters (Figure 2E).

**Figure 2.**
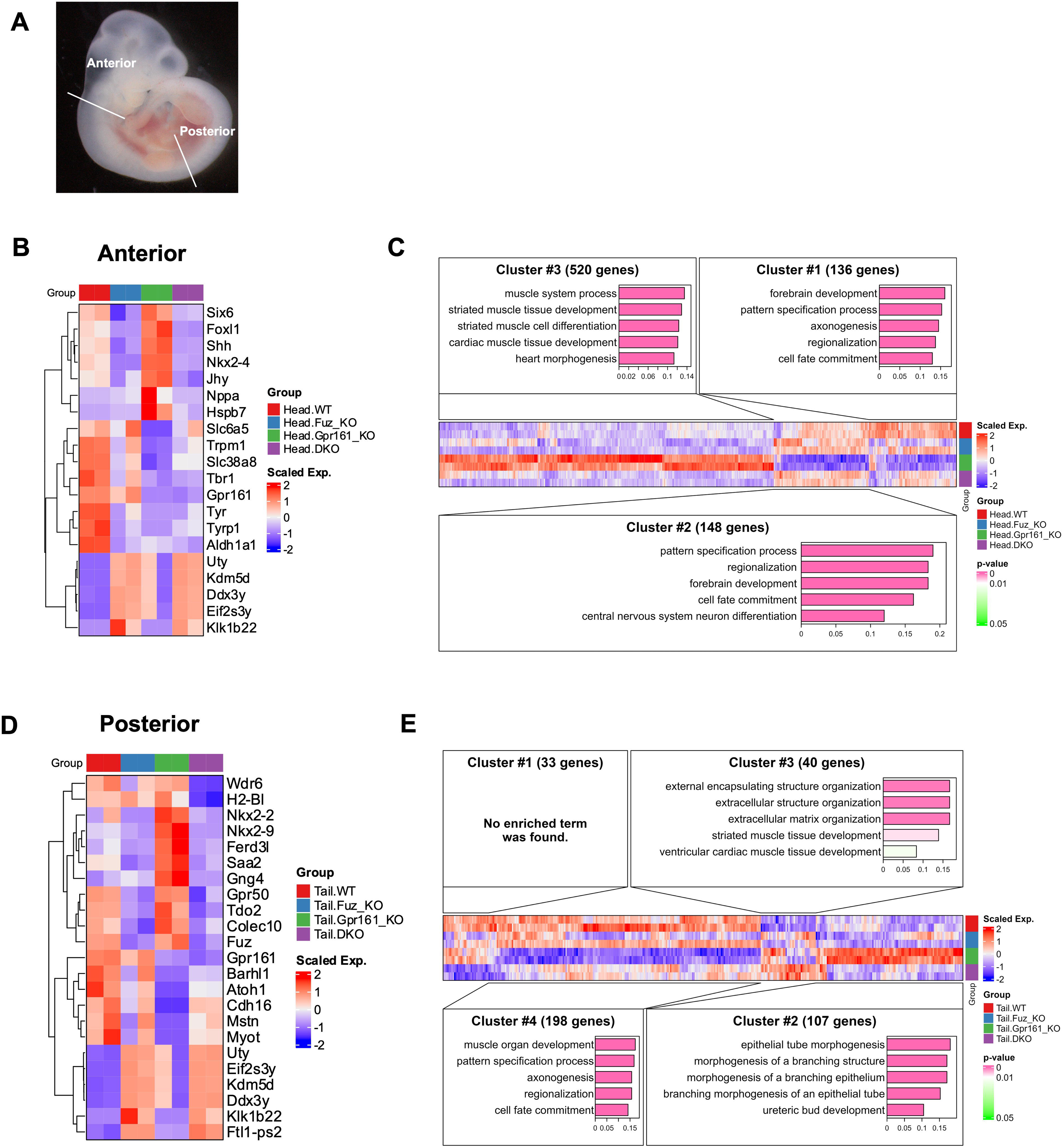
The transcriptomic analysis of anterior and posterior regions of WT, *Fuz* KO, *Gpr161* KO and DKO embryos at E10.5. (A) The schemes of RNA preparations for RNA sequencing. The anterior regions were collected right above the branchial arches, and the posterior regions were collected below the forelimb buds. The anterior and posterior regions of whole embryos were used for RNA preparation. (n=2 for all genotyped embryos) (B) The heatmap for the top 20 differentially regulated genes (DEGs) from anterior regions of each genotyped embryo (WT, *Fuz* KO, *Gpr161* KO and DKO). (C) The gene ontology (GO) analysis with three gene clusters of DEGs from anterior regions of each genotyped embryo. (D) The heatmap for the top 20 DEGs from posterior regions of each genotyped embryo (WT, *Fuz* KO, *Gpr161* KO and DKO). (E) The GO analysis with four clusters of DEGs from posterior regions of each genotyped embryo.

### FUZ biochemically interacted with GPR161

To identify the functional relationship of FUZ and GPR161, we examined whether FUZ and GPR161 physically interacted. We performed co-immunoprecipitation (Co-IP) of FUZ and GPR161 and observed that FUZ interacted with GPR161, confirmed by the reciprocal IP assay (Figures 3A and 3B). The biochemical interaction between GPR161 and FUZ was further validated by a pull-down assay using purified recombinant GST-FUZ proteins. We observed that GPR161 was pulled down with GST-FUZ, but not with GST (Supplementary Figure 3), supporting the interaction between GPR161 and FUZ. To define the domains of FUZ involved in the interaction with GPR161, we performed the immunoprecipitation with C-terminal or N-terminal deletion mutants of FUZ (Fuz dC or FUZ dN). The FUZ dN mutants did not interact with GPR161 (Supplementary Figure 4), confirmed by reciprocal co-immunoprecipitation. This result indicates that N-terminal regions of FUZ are responsible for the interaction with GPR161.

**Figure 3.**
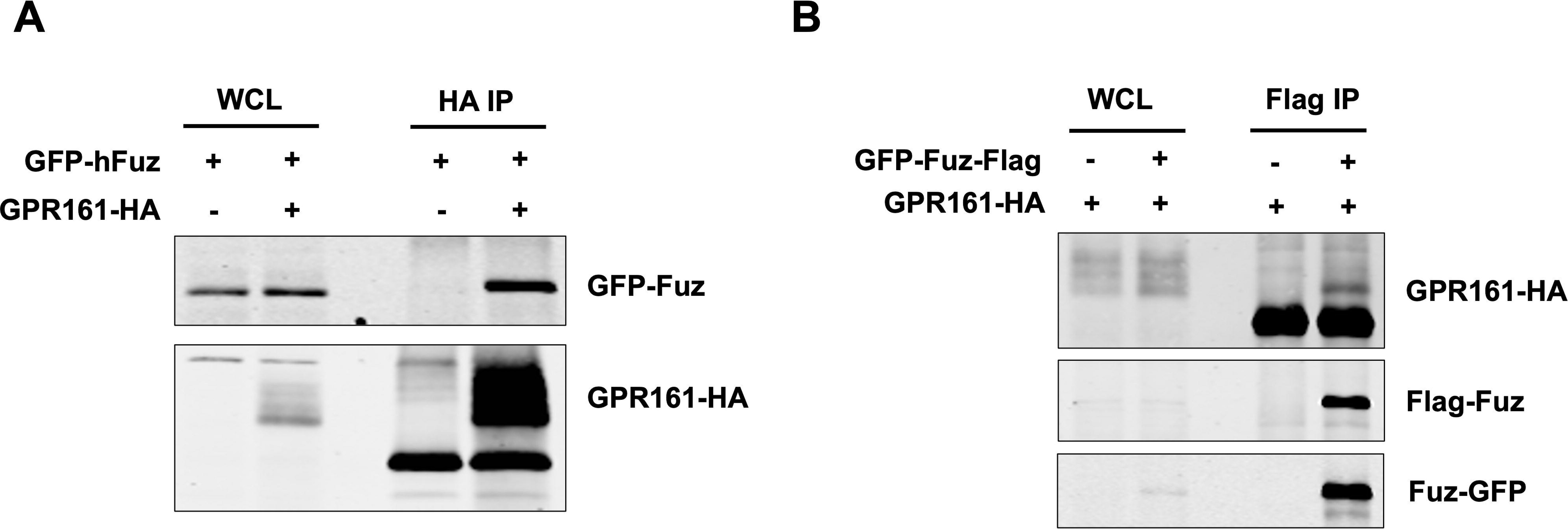
The biochemical interaction between FUZ and GPR161. The GFP-FUZ-Flag and GPR161-HA were co-transfected into HEK293 cells, and the cell lysates were used for immunoprecipitation (IP) with HA antibody (A) and the reciprocal IP with Flag antibody (B). The representative data from three biological replicates.

### FUZ regulated Shh signaling via GPR161 ciliary trafficking

We were interested in understanding how the biochemical interaction between FUZ and GPR161 affected Shh signaling. The mouse phenotypic data suggested that ciliogenesis defects in DKO embryos might not be the only explanation for the genetic relationship between *Fuz* and *Gpr161*. As GPR161 is trafficking in and out of the primary cilia in response to Shh signals, we hypothesized that FUZ affected the ciliary localization of GPR161. The majority of GPR161 (up to 70%) was localized in the primary cilia, and ciliary GPR161 was significantly reduced (∼16%) by the Smoothened agonist (SAG) treatment (Figure 4A). The reduced expression of Fuz by siRNA (up to 85%: Supplementary Figure 5) partially but significantly inhibited the removal of Gpr161 from the primary cilia induced by SAG (Figure 4A). We also observed that the reduced expression of Fuz affected the overall number and length of the primary cilia, as reported in many other studies.

**Figure 4.**
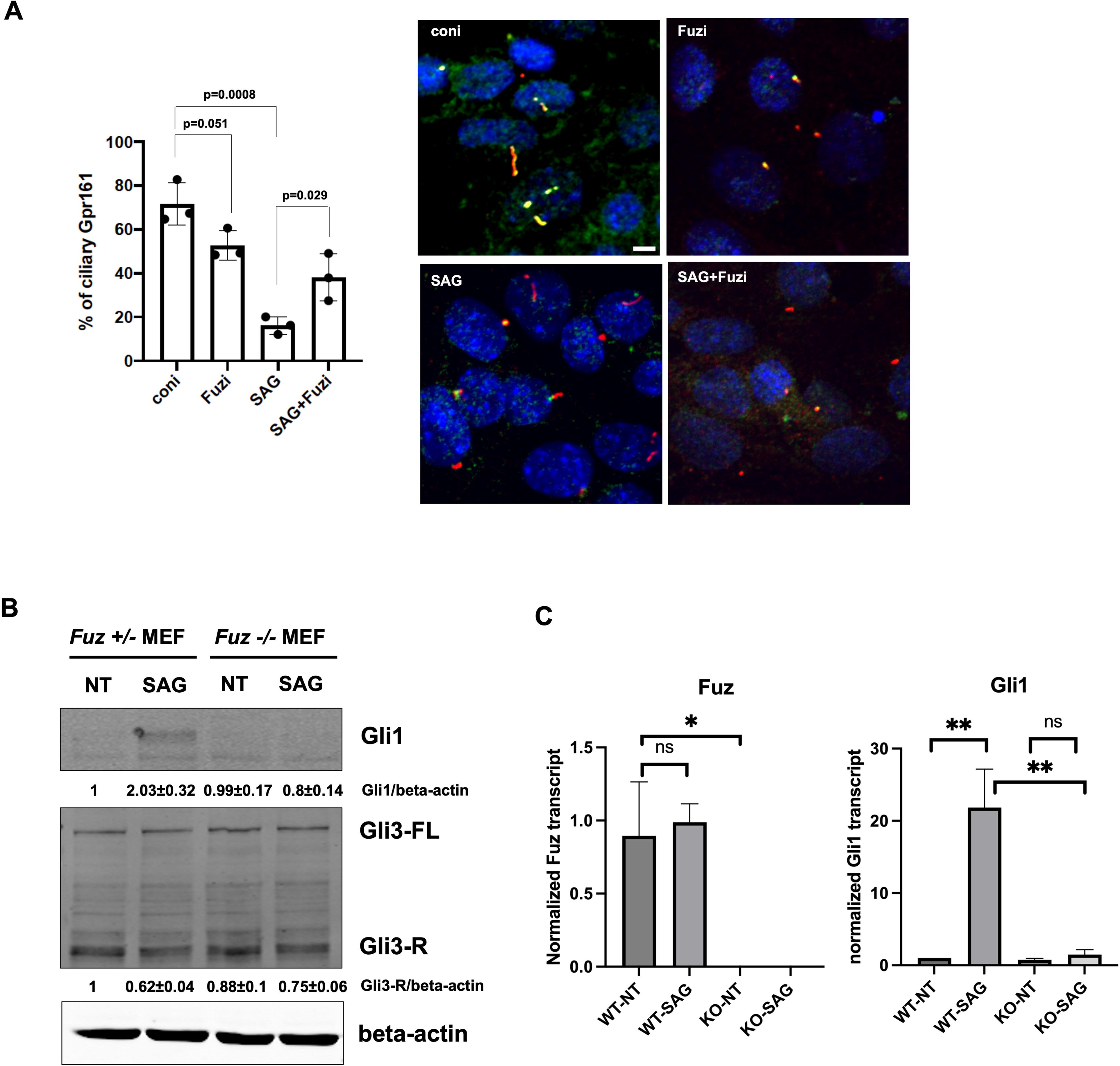
The regulation of Gpr161 ciliary localization by Fuz. (A) IMCD3 cells were treated with Smo agonist (SAG) followed by *Fuz* siRNA transfection for immunostaining with Gpr161 and Arl13b antibodies. n=3, Scale bars: 4 μm. (B and C) The primary WT and *Fuz* KO MEF cells were treated with/without SAG, and the total cell lysates were used for WB with Gli1 (n=3) (B), Gli3 and beta-actin, and for qRT-PCR with *Fuz*, *Gli1* and normalized with *Gapdh* (n=3) (C).

As β-arrestin2 interacts with GPR161 and involves the removal of GPR161 from the primary cilia upon Shh signal (Pal et al., 2016), we evaluated whether FUZ affected the GPR161-β-arrestin2 interaction. The overexpression of FUZ enhanced the biochemical interaction between GPR161 and β-arrestin2 (Figure 5). These results suggested that FUZ might be involved in removing GPR161 from the primary cilia by recruiting β-arrestin2.

**Figure 5.**
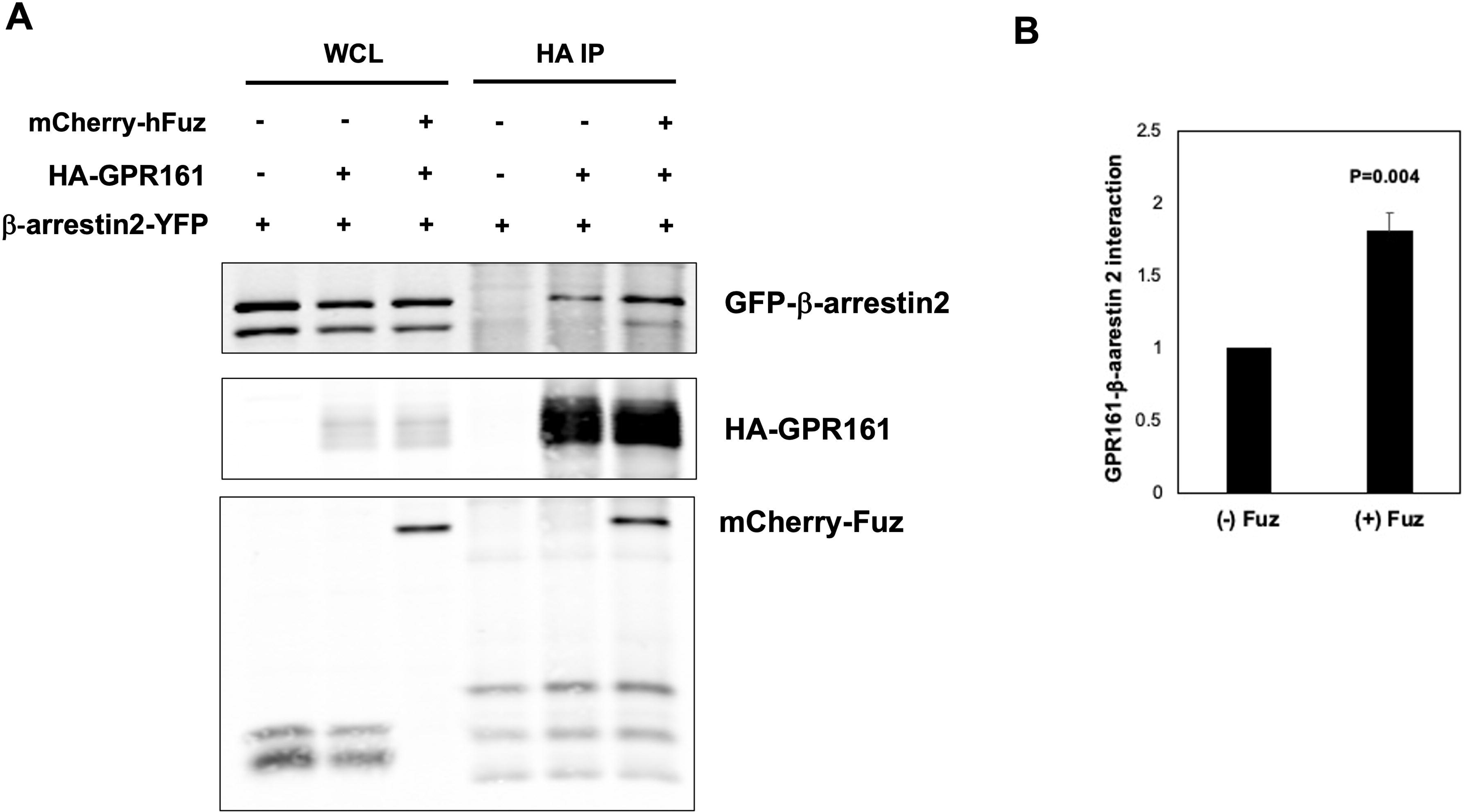
The regulation of GPR161-β-arrestin2 interaction by FUZ. HEK293 cells were transfected with *GPR161-HA* and *β-arrestin2-YFP* with/without *FUZ-mCherry*. The protein interaction was shown by WB followed by immunoprecipitation between GPR161 and β-arrestin2. Right: The quantitation of interaction between GPR161 and β-arrestin2 with/without FUZ (n=3).

We then asked how GPR161 ciliary localization changes by FUZ influenced Shh signaling activities. Both protein and mRNA levels of Gli1 were elevated when SAG was treated in *Fuz+/-* (*Fuz* heterozygote) mouse embryonic fibroblast (MEF) cells, whereas SAG did not significantly change those levels in *Fuz-/-* (*Fuz* KO) MEF cells (Figures 4B and 4C). Consistently, the reduction of Gli3 repressor form, proteolytic forms of Gli3 upon Shh signaling activation, by SAG treatment was more prominent in *Fuz* Het MEF cells than in *Fuz* KO MEF cells. Our results suggested that Fuz involved the removal of GPR161 from the primary cilia via β-arrestin2, thereby regulating Shh signaling.

## Discussion

This study revealed the novel genetic and biochemical linkage between Fuz and Gpr161 in regulating Shh signaling during mouse embryonic development. We determined that Fuz was genetically downstream of Gpr161 during mouse embryonic spinal neural tube formation. We further demonstrated that Fuz regulated Shh signaling via modulating ciliogenesis and the ciliary localization of Gpr161 via β-arrestin 2.

*Fuz* was genetically epistatic to *Gpr161* during the spinal neural tube development since the defects were fully rescued in DKO embryos compared to *Gpr161* KO embryos (Figure 1A). Similarly, cranial vault defects in *Gpr161* cKO with *Prx1-Cre* fetuses were partly rescued with *Gpr161:Ift88* double cKO with *Prx1-Cre* fetuses, supporting Gpr161 mediated intramembranous ossification in mouse cranial vault is partly dependent on the primary cilia (Hwang et al., 2018). Both results support the notion that primary cilia are required for *Gpr161*-mediated spinal neural tube and skeletal morphogenesis during mouse embryonic development. This is a reasonable assumption, given that the inhibitory functions of Gpr161 on Shh signaling largely depend on its localization within the primary cilia. Although the spinal NTDs in *Gpr161* KO embryos were rescued in DKO embryos, the craniofacial defects and cranial NTDs in *Gpr161* KO embryos were partially rescued in DKO embryos, suggesting that Gpr161-regulated craniofacial and cranial neural tube development is not solely dependent on either Fuz or ciliogenesis. Indeed, the comparison between the anterior and posterior regions of WT embryos (data not shown) suggested that cilia-related genes are more abundantly expressed in the anterior regions of embryos. Therefore, other cilia genes might compensate for ciliogenesis activities in the anterior regions over *Fuz* null mutation. This could be one explanation for the partial rescue in the anterior regions of DKO embryos compared to *Gpr161* KO embryos. Furthermore, these data imply that controlling ciliogenesis by genetic modifications or inhibitors may reduce the prevalence or severity of the birth defects, including spinal NTDs, resulting from *Gpr161* null mutants in mice.

Furthermore, the regulatory role of Fuz on Shh signaling might be limited, as we detected no significant changes in Gli1 RNA and protein levels in *Fuz* KO embryos (Figures 1C and 1D). *Fuz* deletion was only functional on Shh signaling when Shh signaling was elevated when *Gpr161* was deleted in mouse embryos shown in DKO embryos. Similarly, the Gli1 RNA and protein levels were also unchanged in *Fuz* KO MEF cells without SAG treatment, whereas those were significantly reduced in *Fuz* KO MEF cells with SAG treatment compared to those in *Fuz* heterozygous MEF cells. These data indicate that Fuz may limitedly act on Shh signaling in certain physiological circumstances or with specific regulators in a tissue-specific manner.

Our RNA-seq data set supported this notion, as the transcriptomic data obtained from the anterior and posterior regions are substantially segregated. In addition, there are only limited numbers (anterior: 53 DEGs, posterior: 22 DEGs) of DEGs between WT and *Fuz* KO embryos compared to those between *Gpr161* KO and DKO embryos for both anterior and posterior regions, consistent with the hypothesis that Fuz may act with Gpr161 in regulating gene expression via signaling pathways, such as Shh signaling during mouse embryonic development. Multiple Shh target genes or related genes were included in the top 20 DEGs from both the anterior and posterior regions of the examined embryos, but there were no overlapping genes except *Gpr16*1 and *Fuz*. Among the top 20 DEGs from the posterior regions of the embryos, the expression of *Atoh1* was decreased in *Gpr161* KO embryos compared to WT embryos, and it was restored in DKO embryos compared to *Gpr161* KO embryos (Figure 2E). Considering the phenotypic rescue of open posterior neuropore (PNP) in DKO embryos compared to *Gpr161* KO, Atoh1 might be the critical transcription factor relevant to mouse spinal neural tube closure regulated by the Gpr161-Fuz axis, which will be an interesting follow-up study.

In the present study, we elucidated the novel biochemical interaction between FUZ and GPR161 and showed that FUZ was involved in GPR161 ciliary trafficking. There were two possibilities concerning GPR161 ciliary trafficking by FUZ: One possibility could be the recruitment of GPR161 to the cilia, and the other could be the removal of cilia. Tulp3 and the IFT-A complex mediate the ciliary recruitment of Gpr161 into the primary cilia (Hirano et al., 2017). The knockdown of Fuz reduced the Gpr161 ciliary localization slightly, but not significantly (Figure 4A), suggesting this possibility is less likely. Instead, the SAG-mediated Gpr161 ciliary removal was significantly reduced when Fuz expression was reduced by siRNA (Figure 4A), indicating that Fuz might affect the removal of Gpr161 from cilia. This possibility was further confirmed by increased interaction between GPR161-β-arrestin2 when FUZ was overexpressed (Figure 5). As Fuz involves retrograde ciliary trafficking, Fuz may regulate the removal of ciliary Gpr161 when the Shh signaling is activated. The involvement of Smo in the Fuz-mediated ciliary removal of Gpr161 was not explored, but it is possible as SAG affected Fuz-mediated Gpr161 ciliary localization. The previous studies suggested that Bardet-Biedl syndrome (BBS) and IFT systems involve the ciliary exit of GPR161 (Ye et al., 2018), and β-arrestin2 might link GPR161 and BBSome via ubiquitination of GPR161 for the ciliary exit (Shinde et al., 2020). One possible mechanism is that FUZ might act to link GPR161 to β-arrestin2, thereby undergoing the ciliary exit by BBSome. We have not tested whether FUZ biochemically interacted with either β-arrestin2 or BBSome, and it will be tested in a future study.

In conclusion, we identified the novel Gpr161-Fuz axis to regulate Shh signaling during mouse embryonic development. As Fuz is one of the effector proteins in PCP signaling, this study shed light on the idea that there might be a cross-regulation between PCP and Shh signaling pathways in not only ciliogenesis/ciliary trafficking but also Shh signaling regulation in multiple developmental contexts.

## Materials and Methods

### Transgenic mouse strains and phenotypic observation

All mouse strains were maintained according to the guidelines approved by The University of Texas at Austin’s Institutional Animal Care and Use Committee (IACUC). *Gpr161* knock-out (*Gpr161* KO) mice were graciously provided by Dr. Saikat Mukhopadhyay (UT Southwestern, Dallas), and detailed information concerning the generation of transgenic lines was previously reported (Hwang et al., 2018). *Fuz* KO mice were generated, and the detail was reported in (Gray et al., 2009). The embryos for phenotypic observation and molecular analysis were collected by timed mating. The embryonic day was determined at E0.5 at noon of the day with positive vaginal plugs. The images of gross phenotypes of embryos were captured with an Olympus SZX2-ILLT microscope (Olympus, Tokyo, Japan). The genotypes of the mice and embryos were determined by PCR-based genotyping.

### Tissue processing for cryosection and immunostaining

Embryos were collected at E9.5 from timed mating and processed for cryo-sectioning. In detail, the embryos from the different litters were fixed, incubated in 30% sucrose solution at 4°C until submerged, and mounted in the optimal cutting temperature (OCT) solution. OCT-embedded embryos were cryo-sectioned with 20 μm thickness for immunostaining. The frozen sections were incubated with blocking buffer (10% normal goat serum in PBS) and washed with PBS. They were incubated with primary antibodies (either anti-Nkx6.1 (Developmental studies hybridoma bank; F55A12-concentrate: 1:200) and Arl13b (Proteintech, 17711-1-AP, 1:200) in blocking buffer), subsequently incubated with secondary antibodies, and then with DAPI (1mg/ml) before mounting. Images were captured with a Nikon Ti2E/CSU-W1 spinning disc confocal microscope.

### DNA plasmids

The DNA plasmids for transfection were *GPR161-HA* (Kim et al., 2019), *GFP-FUZ-Flag* (Seo et al., 2011), and *β-arrestin 2-mYFP* (Addgene #36917). *FUZ-mCherry* was generated by sub-cloning with XhoI/BamHI into *mCherry2-N1* backbone vector (Addgene #54517). FUZ deletion mutants (dN and dC) were generated with the site-directed mutagenesis method with the template (*GFP-FUZ-Flag*). GST-FUZ was generated by sub-cloning with BamHI/XhoI into *pGSTag* backbone vector (Addgene #21877).

### RNA preparation and quantitative RT-PCR (qRT-PCR)

The total mRNA was isolated from cells or embryos with a Direct-zol RNA extraction kit (Zymo Research). RNA (0.5 ug-1ug) was reverse transcribed with iScript RT-Supermix (Bio-Rad), and cDNA was used for qRT PCR with SsoAdvanced SYBR Green Supermix (Bio-Rad). The primers for qRT-PCR were as follows: *Fuz* (Forward: 5’-CACTTGGAACTGCGACGCTG-3’, Reverse: 5’-CACGAGATAACAGGCTCTGG-3’), *Gli1* (Forward: 5’-CCAAGCCAACTTTATGTCAGGG-3’; Reverse: 5’-AGCCCGCTTCTTTGTTAATTTGA-3’), *Ptch1* (Forward: 5’-TGGCTCTTGGAGGGCAGAAATTAC-3’, Reverse: 5’-CCTGGGTGGTCTCTCTACTTTGGT-3’), and *Gapdh* (Forward: 5’-GACCTGCCGTCTAGAAAAAC-3’; Reverse: 5’-CTGTAGCCAAATTCGTTGTC-3’).

### RNA sequencing and data analysis

The quantity and integrity of isolated total RNAs from dissected midbrain tissue were analyzed by Nanodrop (Thermofisher) and Bioanalyzer (Agilent Technologies). The library was prepared with NEBNext Ultra RNA with Poly-A selection and was sequenced on an Illumina Hi-Seq 4000 (Admera Health LLC). Raw sequence reads produced by the sequencer were cleaned using Trimmomatic (version v0.38) (Bolger et al., 2014). The cleaned reads were aligned on the mouse reference genome (GRCm39/mm39) with STAR Aligner version 2.7.1a (Dobin et al., 2013). The featureCounts (version 1.6.0) (Liao et al., 2014) was used to count mapped reads for genes. Differentially expressed genes (DEGs) were identified using DESeq2 version 1.40.2 (Love et al., 2014) with the Benjamini-Hochberg adjusted p-value < 0.05 and fold change > 1.5 threshold for significance. Gene ontology (GO) terms enriched on significant DEGs were accessed using clusterProfiler version 4.9.0.2 (Wu et al., 2021). Heatmaps were generated using ComplexHeatmap (version 2.16.0). All statistical analyses and visualizations were performed under R (version 4.3.2) and the R studio environment.

### Cell culture, transfection, and immunocytochemistry

The mIMCD-3 cells (ATCC-CRL-2123) and HEK-293 cells (ATCC-CRL-1573) were cultured with DMEM/F-12 and DMEM, respectively, with 1% PS and 10% Fetal Bovine Serum (FBS) at 37°C and 5% CO_2_. The primary mouse embryonic fibroblast (MEF) cells were isolated from mouse fetuses except for heads and limbs (*Fuz +/-* and *Fuz -/-* at E13.5) and were cultured with DMEM with 1% PS, 1% MEM, and 10% FBS. For overexpression studies, HEK-293 cells were transfected with respective DNA plasmids and Lipofectamine 2000 (Life Technologies) for 48 hours before lysis. For knockdown studies, mIMCD-3 cells were transfected with *Fuz* siRNA (Dharmacon, On-target plus mouse *Fuz* (70300) J-058818-09, -11) and Lipofectamine RNAiMAX for 48 hours before immunocytochemistry. For SAG treatment, the cells were starved for 24 hours with 2% FBS-containing culture media and were treated with SAG (Sigma #566661: 500 nM) in 2% FBS-containing culture media for another 24 hours before harvest. For immunocytochemistry, cells were fixed with 4% PFA in PBS. They were then incubated with blocking buffer (10% normal goat serum, 1% Triton X-100 in PBS) before primary antibodies (Arl13b), secondary antibodies incubation, DAPI (1 ug/ml) staining and mounting. Images were captured with a Nikon Ti2E/CSU-W1 spinning disc confocal microscope.

### Co-Immunoprecipitation

The HEK-293 cells were transfected, and transfected cells were lysed with the digitonin lysis buffer described in (Pal et al., 2015). The cell lysates were immunoprecipitated with HA (Santa Cruz sc-7392; 1 ug antibodies/500 ug lysates) with protein G Sepharose beads (Thermo Fisher Scientific) or Flag M2 magnetic beads (Sigma M8823) at 4°C overnight, followed by washing with GPCR immunoprecipitation (GPCR IP) buffer. The immunoprecipitated lysates were subjected to WB for further analysis.

### Protein purification and Pull-down assay

GSTag or GSTag-FUZ were transformed into BL21 competent cells (NEB) and the bacteria cells were treated with IPTG for protein induction for 6 hours at 30°C. Cells were lysed with sonication in the lysis buffer (PBS+0.1% Triton X-100+1mM DTT) with lysozyme (1mg/ml) on ice before centrifugation. The soluble fraction was collected for incubation with Glutathione Sepharose beads (Thermo Fisher Scientific) at 4°C overnight before extensive washing with lysis buffer without lysozyme. The GST-bound beads were incubated overnight with the whole cell lysates at 4°C and were followed by extensive washing with GPCR IP buffer. The whole cell lysates were prepared from 293 cells overexpressing GPR161-HA with GPCR IP buffer. The pull-down lysates were subjected with WB.

### Western Blot

The mouse embryos were harvested at E9.5 from timed mating, and the whole embryos were used for protein lysates. The mouse embryos and MEF cells were lysed with Radioimmunoprecipitation assay (RIPA) buffer. The total cell lysates were used for WB analysis. The primary antibodies used were as follows: Gli3 (R&D systems AF3690; 1:1000), Gli1 (Cell signaling #2534; 1:500), GAPDH (Cell signaling #2118; 1:5000), GFP (Cell signaling #2956; 1:1000), HA (Cell signaling #3724; 1:1000), Flag (Sigma F1804; 1:1000 or Cell signaling #14793; 1:1000), b-actin (Cell signaling #4970; 1:5000), and RFP (MBL PM005; 1:3000). The secondary antibodies used were as follows: IRDye® 800CW goat anti-rabbit IgG and IRDye® 680RD goat anti-mouse IgG (LiCOR). The images were captured by Odyssey® (LI-COR). The band intensity was quantified with Image J software (NIH).

### Statistical analysis

The experiments were done in triplicate unless specifically stated otherwise and the data was analyzed by the Standard Deviation (SD) with student t-test for comparing groups (GraphPad Prism10).

## Author contribution

SEK and RHF conceived the overall study, SEK and BJW performed overall experiments, and HYK analyzed the RNA-seq data set. SEK wrote the manuscript draft, and all authors reviewed and edited the manuscript.

## Acknowledgments

We acknowledge Dr. Saikat Mukhopadhyay (UT Southwestern, Dallas, TX) for providing *Gpr161* KO mice and the Histology Core of Dell Pediatric Research Institute for technical assistance for cryo-sectioning.

